# Swarming bacteria undergo localized dynamic phase transition to form stress-induced biofilms

**DOI:** 10.1101/2020.08.11.243733

**Authors:** Iago Grobas, Marco Polin, Munehiro Asally

**Author notes:** Equal contribution (Marco Polin, Munehiro Asally).

## Abstract

Self-organized multi-cellular behaviors enable cells to adapt and tolerate stressors to a greater degree than isolated cells. However, whether and how cellular communities alter their collective behaviors adaptively upon exposure to stress is largely unclear. Here we address this question using *Bacillus subtilis*, a model system for bacterial multicellularity. We discover that, upon exposure to a spatial gradient of kanamycin, swarming bacteria activate matrix genes and transit to biofilms. The initial stage of this transition is underpinned by a stress-induced multi-layer formation, emerging from a biophysical mechanism reminiscent to motility-induced phase separation (MIPS). The physical nature of the process suggests that stressors which suppress the expansion of swarms would induce biofilm formation. Indeed, a simple physical barrier also induces a swarm-to-biofilm transition. Based on the gained insight, we propose a promising strategy of antibiotic treatment to effectively inhibit the transition from swarms to biofilms by targeting the localized phase transition.

## Introduction

The ability to sense, respond and adapt to varieties of chemical, physical and environmental stresses is fundamental to the survival of organisms. In addition to general stress-response pathways at individual cell level, which activate target genes in response to a variety of stresses, multi-cellular systems can tolerate stresses through self-organization, the emergence of order in space and time resulting from local interactions between individual cells (Karsenti, 2008; Wedlich-Söldner and Betz, 2018). Bacterial biofilm formation and swarming are ancient forms of multicellular adaptation (De la Fuente-Núñez et al., 2013; Lyons and Kolter, 2015), where cells can coordinate their behaviors through chemical (Daniels et al., 2004; Xavier, 2011), mechanical (Be’er and Ariel, 2019; Mazza, 2016) and bioelectrical (Benarroch and Asally, 2020; Prindle et al., 2015) interactions. Biofilm cells are much more tolerant to various stresses than the genetically identical cells in isolate (Meredith et al., 2015), owing to the physicochemical properties of extracellular polysubstance (EPS) (Flemming and Wingender, 2010), metabolic coordination (Liu et al., 2015), slow cell growth (Costerton, 1999), and in the case of air-exposed biofilms, diffusion barrier by archetypical wrinkled morphology (Epstein et al., 2011; Vlamakis et al., 2013). Swarming bacteria can also collectively tolerate the antibiotic treatments that are lethal to individual cells-albeit only to a lesser degree than biofilms-through motility-induced mixing and reduced small-molecule uptake (Bhattacharyya et al., 2020; Butler et al., 2010; Kearns, 2010; Lai et al., 2009).

Swarming is a rapid mode of surface colonization (Kearns, 2010), and therefore its ability to withstand high antibiotic concentrations could lead to the subsequent establishment of highly resilient biofilms in regions that could not otherwise have been formed. The formation of biofilms is linked to general stress response pathways (Lories et al., 2020; Nadezhdin et al., 2020) and can be induced, from planktonic cells, by a wide range of biochemical and mechanical stressors, such as aminoglycoside antibiotics (Hoffman et al., 2005), redox-active compounds (Wang et al., 2011), nutrient depletion (Zhang et al., 2014) and mechanical stress (Chu et al., 2018). However, whether swarming collectives may transit into more resilient biofilms upon exposure to stressors, and if so, how such a transition can be initiated within a self-organized swarm, are unknown.

Biophysics of collective motion in bacteria, such as flocking and swarming, is a major topic within the rapidly growing research area of active matter (Bechinger et al., 2016; Geyer et al., 2019; Vicsek et al., 1995). Theoretical models of active matter have repeatedly predicted that, at a sufficient concentration, a collection of motile particles can spontaneously form high-density clusters of particles (Barré et al., 2015; Gonnella et al., 2015). Indeed, this process has been implicated in the development of cell inhomogeneities leading to fruiting-body formation in *M. xanthus* (Liu et al., 2019). This transition, known as Motility Induced Phase Separation (MIPS), is based on feedback between the decrease in particles’ speed at high concentration, caused by physical interactions, and the spontaneous accumulation of active particles in the places where their speed is lower (Gonnella et al., 2015). When the particles’ speed is sufficiently high and their concentration is in the appropriate range (typical volume fractions of 0.3-0.8 or 0.6-0.7 for round or rod-shaped particles, respectively (van Damme et al., 2019; Digregorio et al., 2018)), inherent density fluctuations are amplified by the particles’ slowing down, and the system effectively phase separates into high-density/low-motility clusters surrounded by a low-density/high-motility phase (Cates and Tailleur, 2015).

The theory of MIPS, then, suggests that persistent heterogeneity in cell density-the MIPS clusters-can develop spontaneously when both cell speed and density are appropriate (Fig. 1, grey U-shape region) (Cates et al., 2010). Such conditions should be achievable within a bacterial swarm. Given that cell-density heterogeneity can lead to the production of matrix and biofilm formation mediated by localized cell death (Asally et al., 2012; Ghosh et al., 2013), we hypothesized that the heterogeneity caused by putative MIPS-like clusters could in turn underpin the transition from bacterial swarms into biofilms (Fig. 1). As MIPS-like clustering is an emergent phenomenon arising from physical interactions between individual agents, it may endow swarms with a collective response to a wide spectrum of stressors that cause changes in cell motility and/or density.

**Figure 1.**
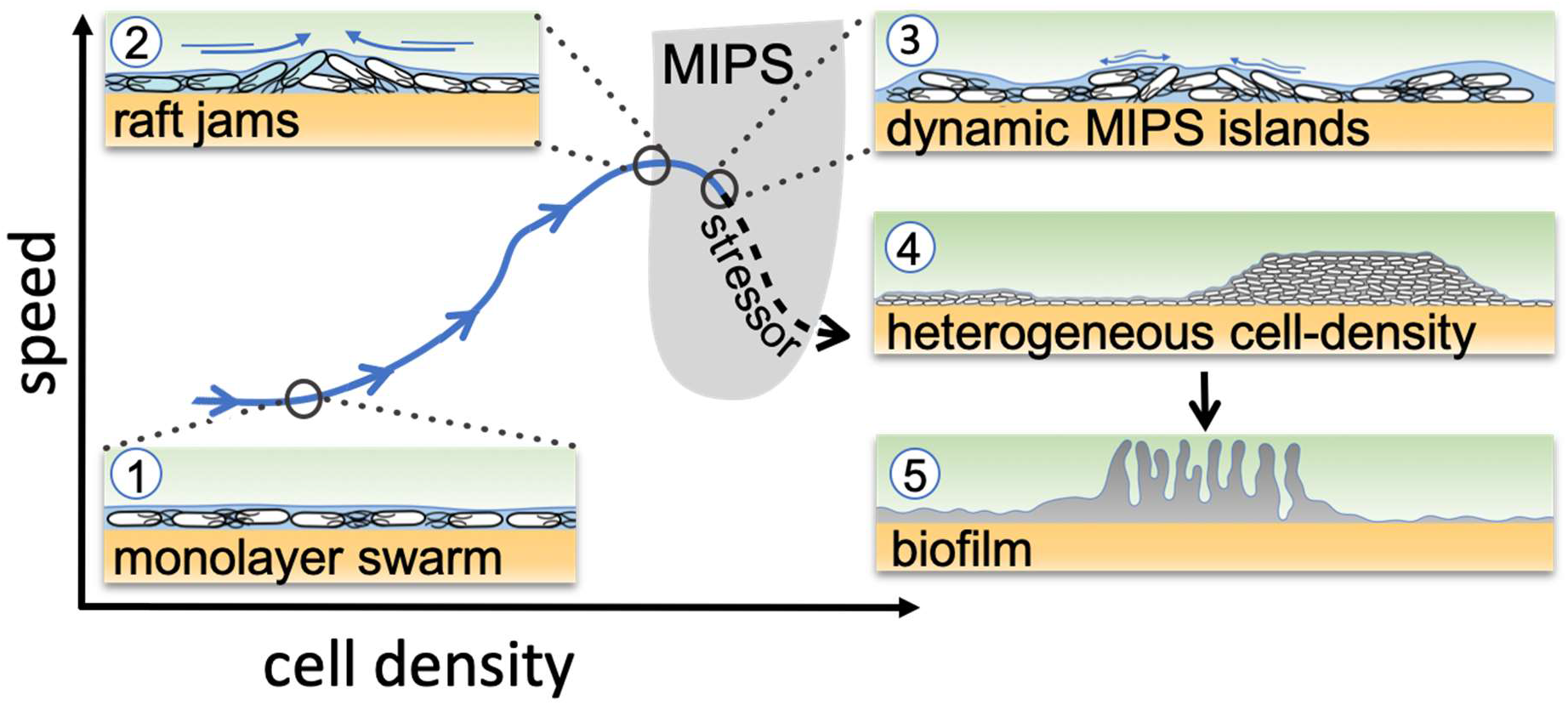
Schematic of the transition from swarming to biofilm formation through MIPS. *B. subtilis* cells swarm in a monolayer (stage 1). As cell density increases, cells form fast-moving rafts which can collide and form transient jams. Cells at the boundary between the colliding rafts are pushed upwards and protrude from the surrounding monolayer (stage 2). Further increase in surface coverage can promote the formation of dynamic MIPS-like islands where cells accumulate within the swarm while still being dynamic (stage 3). Eventually, this uneven distribution of cell density gives rise to macroscopic spatial cell-density heterogeneity in cell density (stage 4), which can lead to the formation of biofilms (stage 5).

Here we show that *Bacillus subtilis* swarms can indeed transit into biofilms through a MIPS-like process, induced by physical or chemical stresses applied at the swarming front. When swarming cells are exposed to kanamycin, they activate the expression of the biofilm matrix operon, *tapA-sipW-tasA*, and eventually develop wrinkled biofilms. The transition initiates by a localized phase transition where the expanding swarming monolayer generates multilayer clusters of cells as a consequence of motility and stress-induced elevation of cell density. Based on the insights gained from our investigation, we show that targeting the multi-layered region by administering a given amount of antibiotic in two separate doses is effective in suppressing the formation of biofilms from swarming cells.

## Results

### Swarming *B. subtilis* transits into a biofilm in presence of a spatial kanamycin gradient

To examine if a stressor triggers biofilm formation from a swarming collective, we performed swarming assays using *B. subtilis* with a spatial gradient of the aminoglycoside kanamycin. Specifically, a disk containing 30 μg of kanamycin was placed on the side of a swarming plate (0.5% agar) and allowed to rest for 24 hours to establish a space-dependent antibiotic concentration profile (Fig. 2a and S1). An inoculum of *B. subtilis* culture was then placed at the center of the dish, ~4 cm away from the kanamycin source, and the plate was imaged while being kept at 30°C (Fig. 2a). After ~2 hours of lag period, the cells formed a rapidly expanding swarming front (~4 mm/h), which became progressively slower towards the source of kanamycin until it completely stopped ~0.3 cm away from the disk (Fig. S2 and Supplementary Video 1). After further incubation at 30°C for 36 hours, the colony developed prominent wrinkles, the morphological hallmark of pellicles and colony biofilms (Cairns et al., 2014; Vlamakis et al., 2013), across a ~3 mm band ~1.3 cm away from the kanamycin disk (Fig. 2b). The estimated level of kanamycin at the site was nearly half the minimum inhibitory concentration (MIC) (Fig. S1). To quantify the degree of biofilm formation, we measured the characteristic wavelength and roughness of the wrinkles, which have been reported to correlate with biofilm stiffness (Asally et al., 2012; Kesel et al., 2016; Yan et al., 2019). The wrinkles appearing near the kanamycin disk had a wavelength of λ = 560 μm and roughness r = 10, while those without kanamycin only produced a faint small-wavelength surface roughness (λ = 91 μm, r = 5.5; Fig. S3).

**Figure 2.**
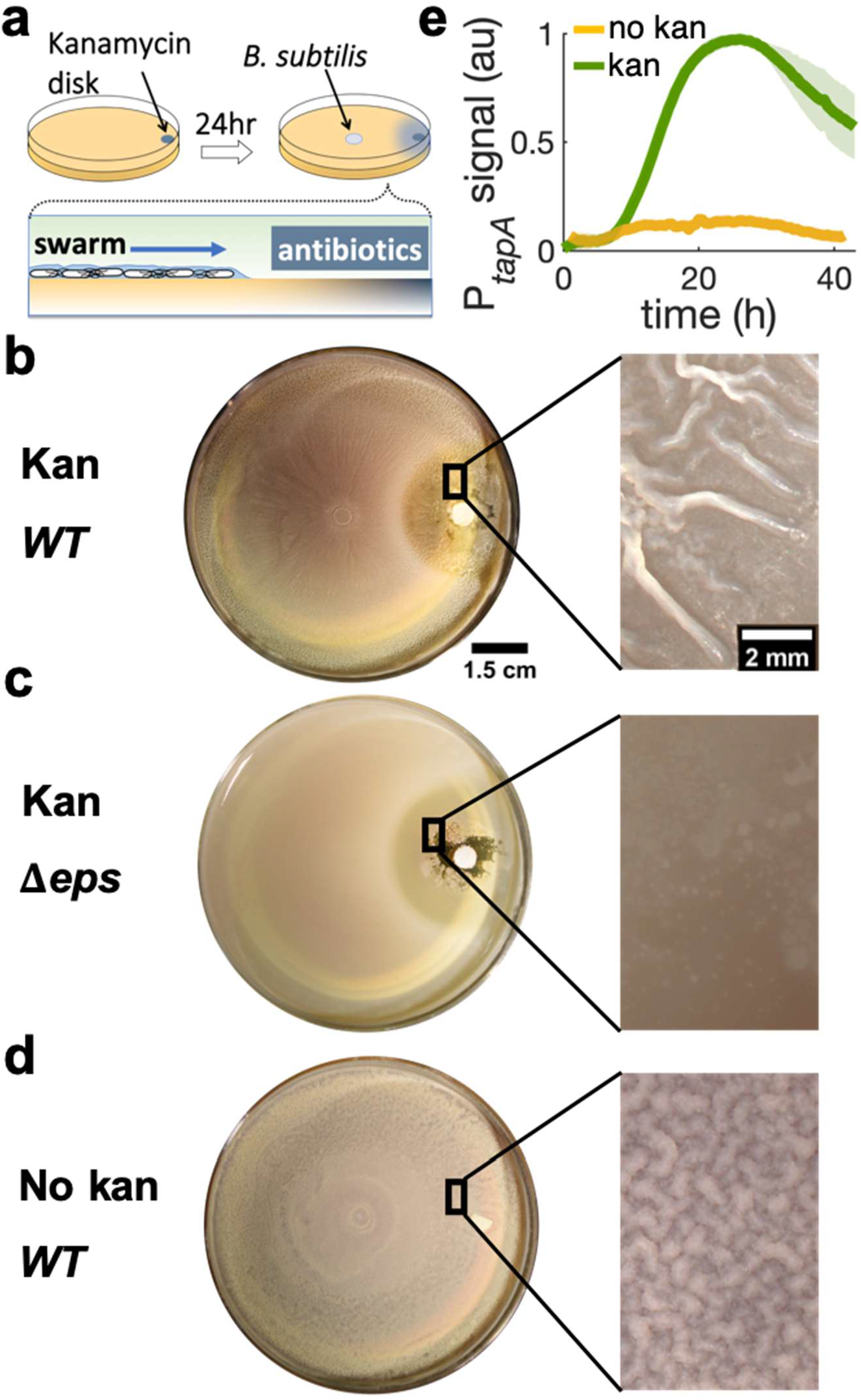
Swarming cells transit into biofilm in presence of kanamycin gradient. a) Schematics of swarming bacteria expanding from the center of a 9 cm Petri dish towards a kanamycin diffusive disk. Kanamycin disk was placed for 24 hours to form a spatial gradient (Fig. S1). b, c, d) Swarming *B. subtilis* plates after 40h incubation. Wrinkles are formed at the region ~2 cm away from disk with b) wildtype (WT), but not with c) *Δ*eps deletion strain. d) WT swarming plate with a diffusive disk without antibiotics. Zoomed images show the colony surface. e) Mean fluorescence intensity of *P_tapA_-yfp* for the front of the plate in presence (green) and absence of kanamycin (yellow). The mean was taken from three independent experiments in the kanamycin case and two independent experiments in absence of kanamycin. The shady areas represent the s.e.m.

To verify the association of these wrinkles to biofilms, we repeated the assay using *Δeps*, a mutant known to be impaired in biofilm formation (Nagórska et al., 2010) but capable of swarming (Supplementary Video 2), and confirmed no wrinkles with this mutant (Fig. 2c). Furthermore, we measured the expression of TasA, an essential matrix component for *B. subtilis* biofilm formation (Romero et al., 2010), using a strain carrying P_*tapA-yfp*_, a fluorescence reporter for the expression of *tapA-sipW-tasA* operon (Vlamakis et al., 2008). The result showed an upregulation of the promoter activity when swarming cells are exposed to kanamycin (Fig. 2e), indicating that the kanamycin activates the matrix gene and induce biofilm formation in a swarming colony on a soft-agar plate.

The emergence of wrinkles across a ~3 mm band away from kanamycin suggests that the swarm-to-biofilm transition corresponds to exposure to a certain concentration range of kanamycin. Indeed, increasing the initial concentration of antibiotic in the disk, wrinkles emerged further away from it (Fig. S4), while a disk without antibiotic did not promote wrinkle formation (Fig. 2d). Intriguingly, on hard agar plates (1.5%) typically used for biofilm assays, wrinkles were only induced to a lesser extent than in the swarming case (Fig. S5). The fact that a kanamycin gradient promotes wrinkle formation in a swarming colony, but not in a non-motile culture, pointed to a fundamental role played by cell motility and suggested the need to investigate the swarming dynamics in detail.

### The emergence of the biofilm is templated by a transition from mono-to multi-layers localized at swarming front

To investigate the potential link between swarming dynamics and biofilm formation, we characterized the swarming dynamics at the single-cell level, with and without a kanamycin gradient. We combined time-lapse imaging at both microscopic (10×; 30 fps) and macroscopic (2×; 0.006 fps) scales to capture both swarming, which is microscopic and fast (~70 μm^2^ cell rafts; speed ~60 μm/s), and biofilm formation, which occurs at a macroscopic scale over hours to days. For the microscopic imaging, we focused on the cells at ~1 cm from the kanamycin disk, where wrinkles eventually appear and kanamycin level is below MIC (Fig. S1). The cells displayed typical swarming dynamics during the expansion of the colony front (Fig. 3a). As time progressed we observed the local surface coverage of the monolayer swarm to progressively increase from initial values of ~20% to ≳60%, at which point the swarming rafts started displaying jamming events lasting ~1-2 sec, during which groups of cells protruded temporarily from the swarming monolayer (typical size of jammed group ~500 μm^2^, see Supplementary Video 3 and Figs. 3a,b).

**Figure 3.**
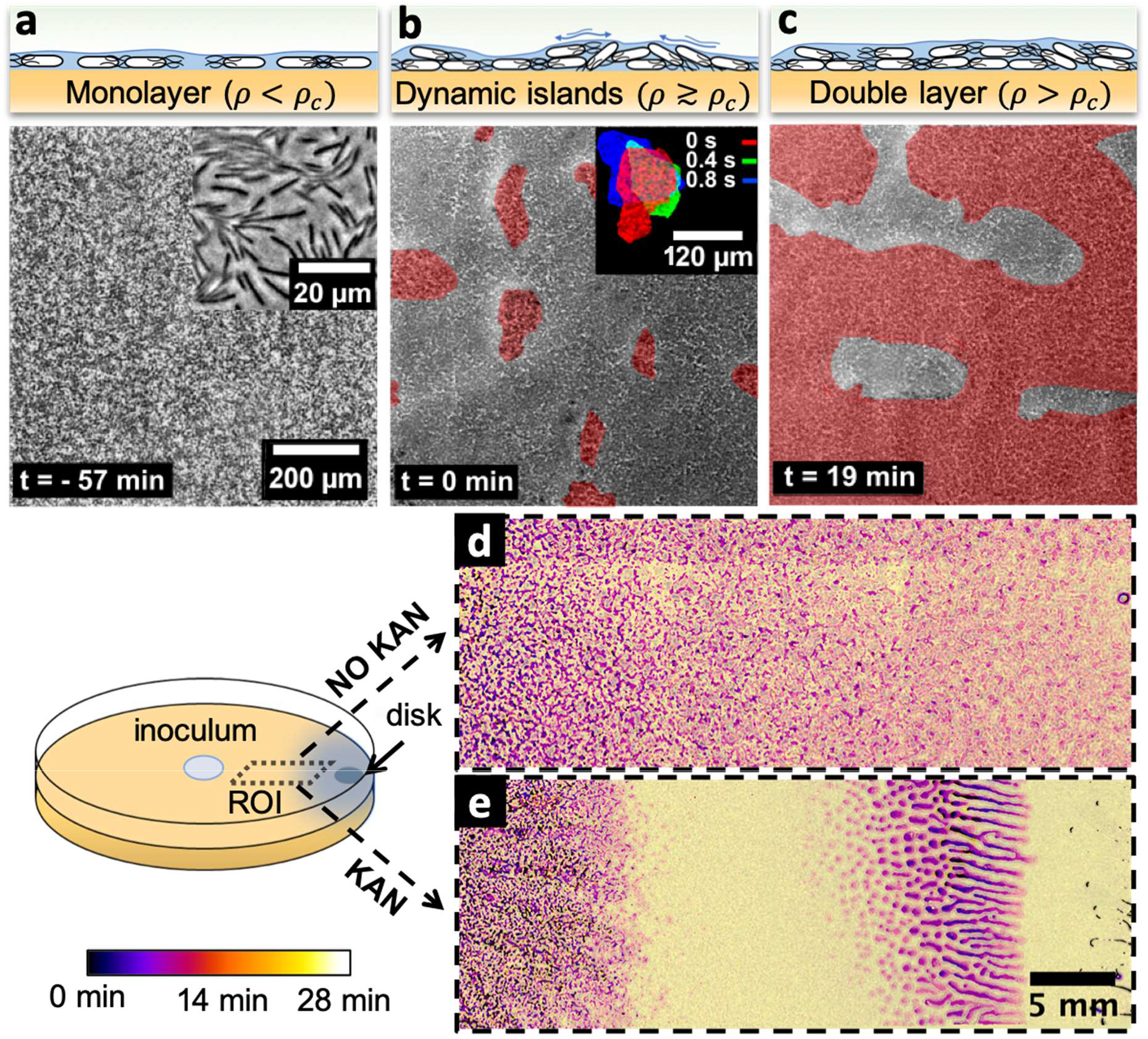
Swarming bacteria form patterned multi-layer regions in presence of kanamycin. a, b, c) Microscopy images of swarming bacteria and side-view illustrations at different levels of cell density (*ρ*). Timestamp is relative to the formation of dynamic multi-layer islands. a) When cell density is below a threshold *ρ_c_*, cells swarm in a monolayer. Image is 57 minutes prior to islands formation. Inset is a zoomed image showing swarming rafts. b) Increase in cell density leads to formation of second layers, highlighted in pseudo red. These islands are highly dynamic, and cells are motile. The shape of the islands changes dynamically within one second (insert). See also Supplementary Video 4. c) Over time, the islands increase their size and merge together to double-layered regions (shown in red), coexisting with mono-layered regions. Double layered regions are seen darker than mono layered regions. d, e) Images of swarming colony with and without kanamycin. Diagram illustrates the regions of interest (ROI) used for panels. The dynamics of island formation are colour coded using the lookup table. The origin of times is the appearance of island. See also Supplementary Video 6. d) In the absence of kanamycin, multilayer regions are much grainer with no clear patterns of propagation. e) In the presence of kanamycin, multi-layered regions have a defined pattern, starting from the regions closer to kanamycin to form an elongated shape. These regions appear predominantly at ~7 mm away from kanamycin.

The jamming of aligned rafts has been predicted numerically for elongated self-propelled particles on a plane (Peruani et al., 2011; Weitz et al., 2015) although in that case the planar confinement precluded any possible excursion in the third dimension. With a slight further increase in local cell density, the temporary jams led to permanent isolated multi-layered regions, which we call ‘islands’, typically ~27,000 μm^2^ in size and with constantly fluctuating boundaries (Fig. 3b inset, Supplementary Video 4). The islands did not appear to be related to any evident inhomogeneity of the underlying agar. Within islands, cells were highly dynamic and appeared to meander across the layers albeit at a reduced speed. Cell movement within islands (Fig. S7, Supplementary Video 5) highlights the fact that these islands are formed by actively motile cells, rather than groups of immotile bacteria which spontaneously separate from the monolayer. In particular, we did not find any evidence for clusters of immotile cells templating the islands. Rather, motile cells were constantly exchanged between the multilayer and the surrounding monolayer. Following a continuous increase in cell density with time, islands grew in size and merged with each other, eventually forming a much larger multi-layered region (Fig. 3c, Fig S7, Supplementary Video 6). The nucleation and merging of islands were observed up to 4 layers before we lost the ability to recognize further transitions due to a flattening of the image contrast (Fig. S7).

The formation of islands was also observed by macroscopic time-lapse imaging. In absence of kanamycin, islands emerged simultaneously throughout the plate, forming an intricate granular pattern with a typical size of ~2,000 μm^2^ (Fig. 3d), a phenomenology reminiscent of spinodal decomposition in binary fluids (Qiu et al., 2001). In contrast, the process of islands formation and growth was markedly different in presence of kanamycin (Fig. 3e). Following the halt of the swarming front expansion due to inhibition by the antibiotic, well-defined, distinct islands emerged well separated and only within a ~5mm-wide band just inside the arrested swarming front, and then grew with a strongly anisotropic pattern oriented transversally to the front of the swarm (Video S6). This phenomenology is reminiscent of a binodal-rather than spinodal-liquid-liquid phase separation in a temperature gradient (Bartolini et al., 2019) which here might reflect a kanamycin-induced speed gradient. The latter macroscopic cell-density heterogeneity resulted in the subsequent formation of biofilm wrinkles, ~1 cm away from the disk (Fig. 2b).

Since quorum sensing is often associated with bacterial collective behavior, we wondered if quorum sensing may play a role in the emergence of islands and the ensuing multilayer. We therefore repeated the experiments with *ΔphrC* and *Δopp* mutant strains, lacking the Phr quorum-sensing system in *B. subtilis* but still capable of swarming. In both cases, we observed the wild-type phenomenology of wrinkled biofilms and the emergence of islands (Fig. S8). The results indicated that this quorum-sensing system is not responsible for the multi-layer transition, suggesting that exposure to kanamycin can promote cell-density heterogeneity by multi-layer islands formation in a quorum-sensing independent manner.

### Transition from monolayer to multilayer resembles motility induced phase separation

To address the mechanism of multi-layer formation, we wondered if physical stresses may be responsible for this emergent collective behavior since single-to-multi layer transitions have been reported for confined bacterial aggregates, either growing or gliding, as a consequence of build-up of internal mechanical stresses (Grant et al., 2014; Su et al., 2012). In particular, we considered if the active matter model of MIPS for rod-shape particles (Cates and Tailleur, 2015) could act as a useful paradigm for understanding the emergence of multi-layer regions. We therefore mapped our experimental results onto the typical phase space considered when studying MIPS transitions, where an active 2-dimensional system is characterized by its surface coverage, φ, and its rotational Péclet number, Per = u/LDr. The latter is defined in terms of the average speed u, characteristic size L, and rotational diffusivity D_r_ of the active particles. MIPS clusters are expected within a U-shaped region characterized by a sufficiently large Per and a range of surface coverages that, for rod-like particles like *B. subtilis*, is pushed to values higher than the ~50%, typical of circular particles, due to antagonistic effects of cell-to-cell alignment (Fig. 4a, grey U-shape region shows the prediction for aspect ratio 2 from (van Damme et al., 2019)). We then quantified the cell density (surface coverage, φ) and the rotational Péclet number from the different stages of the swarming process. Figure 4a shows these two quantities for cells in the monolayer (blue dots) and for bacterial jams (red dots). While bacteria in the monolayer corresponded overwhelmingly to points well outside the MIPS region (Fig. 4a blue dots), bacterial jams-the first stage in the development of stable multilayer islands-clustered around the area predicted for MIPS in two-dimensional self-propelled rods (Fig. 4a, red dots; MIPS region from (Grant et al., 2014)). Jamming events were also characterized by a sudden drop in cells’ speed (Fig. 4b and Supplementary Video 3), consistent with the basic premise of MIPS. Altogether, these results show that the mechanism leading to the development of multilayered islands is compatible with the MIPS process in active matter. Having established the similarities between MIPS and multilayer formation in the swarm, it is important to test the ability of the model to predict the results of new experiments.

**Figure 4.**
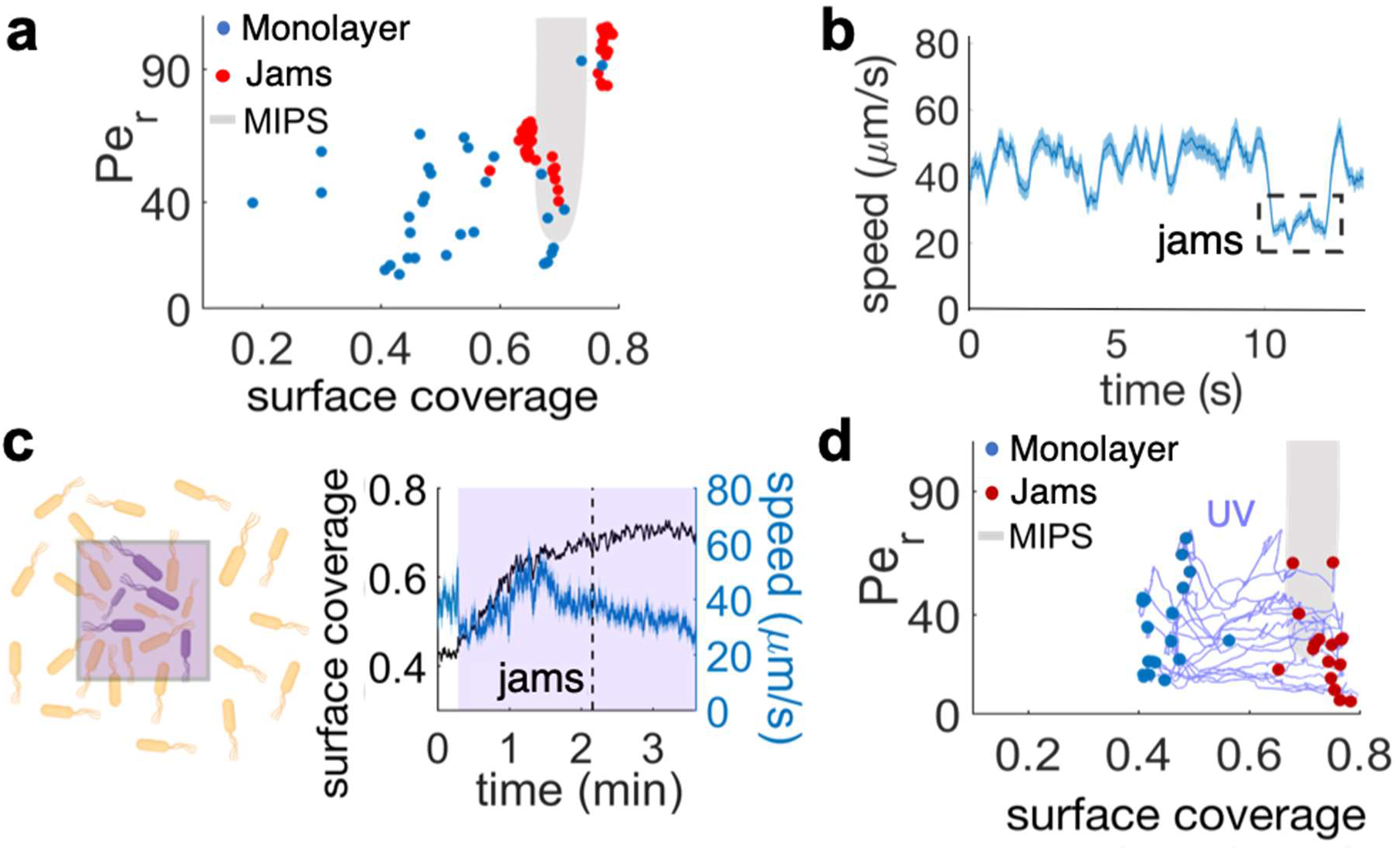
The interplay between cell density and cell speed primes jamming and multi-layer formation. a) Phase diagram of surface coverage and the rotational Peclet number (*P_e_r__*). *P_e_r__* is proportional to the cell motility speed (see Methods). Greyed region depicts motility-induced phase separation (MIPS) heterogeneity. Each data point represents a result of a 4-sec single-cell time-lapse microscopy data. The time-lapse data in which jamming events are observed are shown in red (see Supplementary Video 3). Jams appear exclusively under the condition near the border of MIPS region. b) The speed of motile cells drops by half during jamming (highlighted by dashed rectangle). Increase in jamming events lead to formation of islands (see Fig. 2 and Supplementary Video 4). c, left) Illustrative diagram showing the accumulation of cells by UV irradiation. Cell speeds drop within the irradiated region (show by purple square), which elevates the surface coverage. c, right) UV irradiation elevates surface coverage. Graph shows the dynamics of surface coverage (black) and cell speed (cyan). Magenta is the period with UV light illumination (1.2 mW/mm^2^). Vertical dashed line shows a jamming event. d) Time evolution trajectories of surface coverage and rotational Peclet number with UV illumination experiment. Blue dots are before UV illumination and red dots are when jams appear. Increasing surface coverage by UV induces formation of jams at the border to MIPS region (grey).

### Local accumulation of swarming cells induces multilayer transition and biofilm formation

The MIPS paradigm makes the experimentally testable prediction that it should be possible to induce multilayer formation by altering the local density of motile cells, thereby forcing the system to enter the MIPS region in phase space (Fig. 4a, shaded region). We tested this in two ways: by UV irradiation, and through the use of a physical barrier to block front expansion. Near-UV light can decrease cell speed in gram-negative bacteria like *E. coli* and *S. marcescens* (Krasnopeeva et al., 2019; Yang et al., 2019), as well as, as we report here, in *B. subtilis* (Fig. S9a). Locally slowing down motility by light can lead to a local increase in cell concentration through the accumulation of cells from non-irradiated regions (Fig. 4c). We first verified that the illumination by UV light itself does not induce a multilayer transition, by illuminating an area >200-fold greater than the field of view. This prevented cells outside of the irradiated area from accumulating within the field of view during the experiment. Accordingly, the surface coverage increased only marginally (φ ≃ 0.45 to φ ≃ 0.47 in 3 min; Fig. S9a). The UV illumination caused an initial sudden drop in average speed from 65 μm/s to 50 μm/s (Fig, S9a), followed by a nonlinear progressive slow down over the course of 3 min. Within the phase space picture (Fig. 4a, Fig. S9b), this corresponds to a trajectory that essentially just moves towards progressively lower values of Pe_r_. This should not lead to either jams or island formation, which in fact were never observed (Fig. 4a, Fig. S9b).

We next illuminated a region of size similar to the field of view. This arrangement allowed cells from the outer region to accumulate within the field of view due to UV-induced slow-down, a phenomenon which is a direct consequence of the active, out-of-equilibrium nature of the swarm with no counterpart in statistical systems in equilibrium (Arlt et al., 2018; Cates, 2012; Frangipane et al., 2018). As a consequence, the cell density increased significantly (φ ≃ 0.42 to φ ≃ 0.7 in 2 min; Fig. 4c). For the first ~2 min, the average drop in cell speed was likely compensated by the density increase. It is well known, in fact, that cell density can enhance swarming motility through cooperative raft formation (Be’er and Ariel, 2019; Jeckel et al., 2019). Eventually, however, the increase in cell density resulted first in the formation of jams and finally of multi-layer islands (Fig. 4c, Supplementary Video 7). Figure 4d shows the trajectories followed by the irradiated swarms in phase space. Jamming of swarming cells, the first step in island formation, occurs only for cell densities within a range that compares very well with predictions by MIPS for self-propelled rod-like particles (Fig. 4d shaded region and (van Damme et al., 2019)). These results provide a direct support to the hypothesis that a biophysical MIPS-like process underpins the transition from swarming monolayers to multilayers in *B. subtilis*.

To further examine if a local cell-density increase is sufficient by itself to induce a localized transition from monolayer to multilayer, we used a physical barrier to impede the advance of the swam and locally increase cell density. Again, consistently with the MIPS picture, the arrest of the swarm front led to an increase in cell density and the subsequent emergence of multi-layer islands near the barrier (Supplementary Video 8). After 36 hours of further incubation, wrinkles developed near the physical barrier precisely in the region where the islands had started to appear initially (Fig. 5a).

**Figure 5.**
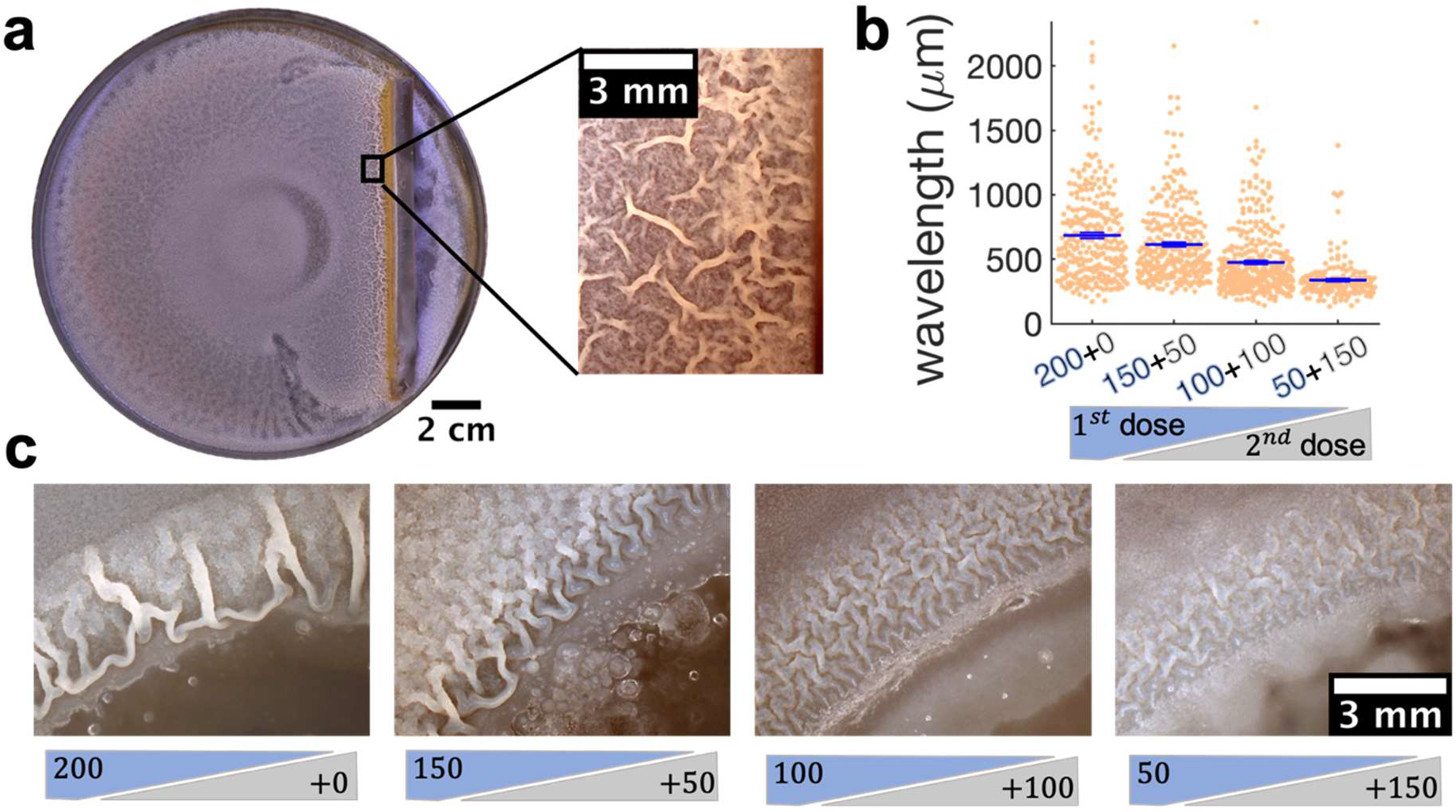
Locally increasing cell density is sufficient in inducing wrinkle formation, which can be inhibited by sequential administration of kanamycin. a) A physical barrier placed on agar triggers wrinkle formation. Wrinkles are formed at the region close to the barrier. A plate was incubated for 40 hours as in Figure 1. b, c) Kanamycin was sequentially administered in two different doses to a disk while keeping the total to be 200 *μg*. The first administration was added as in other experiments, while the second was added to the disk when the islands were about to appear. b) The wavelengths of wrinkles with respect to the doses of first (blue font) and second (gray font) administrations (*μg*). c) Microscopy images of wrinkles induced by kanamycin. The first dose is in the blue shade and the second dose in the grey shade. Even when the total dose is kept the same, equal or greater 2^nd^ dose inhibits biofilm formation.

Altogether, these results strongly support that *B. subtilis* swarms can undergo a single-to-multi-layer transition driven principally by a physical mechanism compatible with MIPS, which can be either global and spinodal-like, or localized and binodal-like. When this transition is localized, regardless of it being caused by antibiotics or physical confinements, the resultant macroscopic cell-density heterogeneity determines the emergence of wrinkled biofilms.

### Sequential administration of antibiotics suppresses the emergence of biofilms from swarms

While more complex signaling pathways regulating biofilm matrix production are likely involved in the passage from localized multi-layers to wrinkles, our results suggest that altering the expansion dynamics of a swarm promotes biofilm formation. This implies that exposing bacterial swarms to stressors, such as kanamycin, physical barrier and UV light, may inadvertently increase their resilience by promoting the formation of biofilms that are much more difficult to eradicate. This is difficult to prevent by simply increasing the amount of antibiotics. In fact, when we used a ~7-fold greater dose of kanamycin in the diffusion disk (200 μg), wrinkles still appeared on the plate, although at a greater distance from the disk (Fig. S4). However, our findings suggest that the multi-layer band could be a good target for antibiotic treatment aimed at suppressing the emergence of biofilms. Such multilayer band happens at a concentration of antibiotic that bacteria can tolerate since the cells are still motile (in our case ~0.5 MIC; see Fig. S6, Supplementary Video 5). We therefore wondered if a two-step sequential antibiotic administration could prevent biofilm formation, where the first administration induces multilayer formation and the second targets the multilayer area before completing biofilm formation. To test this conjecture, we decided to administer a total amount of 200 μg of kanamycin in two steps, an initial one when placing the disk on the plate, and the second as the swarming front stopped (Fig. 5b, Fig. S10). The emergence of wrinkles was greatly suppressed when kanamycin was administrated sequentially, despite keeping the total amount of antibiotic constant (Fig. 5b,c and Fig. S10). The effect was most evident when the second dose was greater than the first. These results thus propose a promising strategy for treating bacterial collectives with aminoglycoside while minimizing the emergence of biofilms.

## Discussion

This work reveals a biophysical mechanism underpinning the initial stage of the collective stress-induced transition from swarms to biofilm in *B. subtilis* governed by motility and cell density. This is based on the halting of swarming expansion, which promotes the accumulation of cells at the front, resulting in the formation of MIPS-like multi-layered islands. Upon exposure to the aminoglycoside kanamycin, the swarming colony activates the *tapA-sipW-tasA* operon and eventually develops a wrinkled biofilm morphology. Consistent with this view, we demonstrate that qualitatively different stressors, from antibiotics to UV and physical confinement, can all induce formation of islands.

Moreover, based on our findings, we show that a sequential monotherapy can be effective in preventing biofilm formation from a swarming colony in *B. subtilis*. As the underpinning mechanism of the transition is an emergent phenomenon driven by physical interactions between swarming cells, we believe similar transitions should also happen in other bacterial species. It would be interesting, for example, to examine if swarms of clinically relevant bacteria, such as *Pseudomonas aeruginosa* and *Salmonella enterica*, may also transit into biofilms through a similar process. Such investigations would be an important step forward to see if sequential administrations could be effective in preventing stress-induced biofilm formation also in pathogenic bacteria.

This study addresses a fundamental question about the mechanism by which cell collectives adapt their behavior in response to various physical and chemical stresses. In the present case, a local cell-density increase caused by the halting of the swarming front, may be part of a general collective stress response mechanism, which triggers a switch in the collective behavior from swarming to biofilm. Such a stress response mechanism at the collective level could allow the swarming colony to develop biofilms in response to various stressors, regardless of the stressors’ exact molecular mode of action. We expect that further research will determine whether this form of environmental sensing and adaptation of cell collectives via cell-density increase is common to other biological systems. Interestingly, this idea is in line with recent advancements in the understanding of mechanochemical feedback in development and disease, where local cell density can determine the fates of cell collectives (Hannezo and Heisenberg, 2019). The connection we discovered could represent a primitive example of a collective mechanochemical feedback loop, underpinned by one of the most fundamental types of emergent phenomena (MIPS) in collections of motile agents either alive or synthetic. To this end, the gained biophysical insights may not only offer new biomedical treatment strategies against the rise of biofilm-associated antimicrobial resistance but may also contribute to our understanding of development and cell-fate determination.

Following our discovery of stress-induced swarm-to-biofilm transition, we present a detailed characterization of the initial stage of the transition, namely development of multilayered islands. As with every discovery, this also brings a host of new questions. For example, it is unclear whether the molecular mechanisms driving biofilm development from planktonic cells is identical to the one from swarms. We show the activation of *tasA* gene during the kanamycin-induced transition from swarms to biofilms, suggesting the commonality between these processes. The expression of biofilm matrix genes is regulated by various complex pathways, where Spo0A, SinR and AbrB being central regulators (Vlamakis et al., 2013). If swarms develop biofilms through different pathways, elucidating the molecular regulatory machinery during the transition from swarm to biofilm may unveil new molecular pathways regulating biofilm formation. From the perspective of biofilm being a multicellular adaptation, it would be interesting to determine the stage at which swarming collective loses the ability to adapt to environmental changes. The biophysical transition that we report here would suggest that this happens once the ageing swarm cannot be driven anymore to within the putative MIPS region by the external stressor. Another important question lies in the interplay between biophysical and molecular mechanisms regulating stress-induced biofilm development. Characterizing the gene expression profiles in the high-cell-density clusters resulting from multilayered islands, while simultaneously monitoring the mechanical interactions, would be an important step forward towards gaining a holistic understanding of collective stress response. We hope that our work will inspire new research in this area, and look forward to further exciting results in the near future.

## Materials and Methods

### Kanamycin gradient assay

−80°C glycerol stock of *Bacillus subtilis* NCIB3610 wild-type strain (WT) was streaked on a lysogeny-broth (LB) 1.5 % agar plate and grown overnight at 37°C. When specified in figure caption, a genetically modified strain (listed in Table 1) was used instead of WT. A single colony was picked from this plate and incubated in 1 ml of liquid LB for 3 h at 37°C. A 4 *μ*l inoculum from this culture was placed in the center of a 0.5% LB agar plate supplemented with 2% of glycerol and 0.1 mM MnSO_4_ (LBGM (Shemesh and Chai, 2013)) to favor biofilm formation. A kanamycin diffusive disk (Oxoid™ 30 *μ*g) was placed on a side of the plate 24 h before inoculation to allow the antibiotic to diffuse at room temperature. The distance between the inoculum and the kanamycin disk was approximately 3.2 cm, and plates were incubated for additional 40 h after inoculation at 30°C.

### Images of swarming plates

Low magnification images of the plates were acquired with a DSLR D5000 Nikon camera (lens AF-S Micro NIKKOR 40 MM 1.28) in a 30°C incubator. The incubator was covered by black tape to avoid reflections and the illumination was provided by a white LED placed on a side of the plate. Higher magnification images (e.g. of the wrinkles) were taken by an Olympus SZ61 microscope by placing the plates in a dark background with illumination coming from a LED ring attached to the microscope.

### Quantification of the P_*tapA*_-*yfp* reporter

Biofilm extracellular matrix production was characterized by using a modified strain carrying P_*tapA*_-*yfp*, a fluorescence reporter for the expression of *tapA-sipW-tasA* operon. The kanamycin gradient assay was repeated by inoculating this strain in LBGM (1.2% and 2% glycerol) swarming agar plates. The experiment was replicated three times with three different lenses and microscopes: 2x (Nikon Apo Lambda 2x UW, NA 0.1), 2x (Nikon Plan 2x UW, NA 0.06), 2.5x (Leica 2.5x N PLAN, NA 0.07); microscopes: two Nikon Eclipse Ti2 and Leica DMi8. Images were taken every half an hour for a period of ~40 h in a region of 3×9 cm^2^ going from the disk to the inoculum. The images were stitched using the “Grid/Collection stitching” plugin (Preibisch et al., 2009) in Fiji (Schindelin et al., 2012). The experiment was repeated twice in absence of kanamycin with the first two microscopes and lenses. To calculate the fluorescent signal, a region of interest was drawn in Fiji surrounding the area where the wrinkles appeared. The signal was normalized by subtracting the minimum value of pixel intensity recorded in the time-lapse and dividing by the maximum of the signal for each of the microscopes.

### Raft and islands sizes

The size of the rafts within the swarm were obtained from three different experiments. A freehand line drawn with Fiji/imageJ (Schindelin et al., 2012) enclosing the raft was used to measure the area within the line. Measurements across different positions and time points were used to account for the variability in raft size within a swarm depending on the position and/or time after expansion begins (Jeckel et al., 2019). To measure the islands size, timelapses of islands formation in presence and absence of kanamycin were used. The first frame (just before islands appeared) was subtracted to the timelapse and then a gaussian filter was applied to remove noise. Finally, the timelapse was threshold and the initial size of the islands was measured by ‘regionprops’ in Matlab.

### Characterization of island formation

We characterized the double layer using 2x (Nikon Plan 2x UW, NA 0.06), 2.5x (Leica 2.5x N PLAN, NA 0.07) and 10x (Nikon 10x PLAN FLUOR PH2 DLL, NA 0.3) magnifications in two different microscopes, Nikon Eclipse Ti2 and Leica DMi8. The images were acquired every 2 min in the Nikon Eclipse Ti2 and every 1 min in the Leica Dmi8 using brightfield illumination. Cell motility and cell density were recorded with a 40x objective (Nikon 40x LWD, NA 0.55) at 37°C. Cell motility was measured by adapting a Particle Image Velocimetry (PIV) code written in Matlab (Sveen, 2004). The surface coverage was measured by thresholding the images and dividing the area covered by bacteria by the total area of our field of view. The threshold was set by using the command ‘imbinarize’ in Matlab and adapting the sensitivity of its threshold to account for the best estimate of cells in the field of view.

### Near UV experiments

Bacteria were irradiated by near UV-Violet light for 30 s using the inverted microscope DMi8 (Leica Microsystems) and the LED light source, SOLA SM II Light Engine (Lumencor), with an excitation filter 400/16 using a 40x objective (Leica 40x PH2 HC PL FLUO, NA 0.6) for approximately 3.5 minutes at an intensity of 1.2 to 5.3 mW/mm^2^. The light intensities were measured by placing a photodiode power sensor (Thorlabs S120C) on the microscope stage. To irradiate a larger area than the one with the epi-fluorescence set up, a Thorlabs 405 nm light LED was coupled to a cage cube (see Table S1 for details about the components of the set up). The light coming from the LED was concentrated by an aspheric condenser lens and reflected by a 45° dichroic mirror towards the sample. The area illuminated is roughly a circle of 2.5 mm radius at an intensity of 1.5 mW/mm^2^. The surface coverage was calculated by binarizing the time lapses applying a locally adaptive threshold in Matlab. The sensitivity of such threshold was changed along the time lapse when cell density was rapidly increasing and the same value of the threshold could not account for the total number of particles.

The trajectories of the swarming bacteria in the phase diagram were calculated by calculating the Pe_*r*_ and the surface coverage as detailed in the previous section. The trajectories are the result of continuous irradiation with either of the setups previously described. When illuminated by the epifluorescence the trajectories were plotted until a jam of bacteria was observed in the field of view. When shining UV from the condenser the trajectories were plotted until bacteria were completely immotile (Fig. S9). Data for speed and surface coverage of bacteria has been smoothed by the ‘smooth’ function in Matlab.

### Quorum sensing experiments

The same protocol as the explained in ‘kanamycin gradient assay’ was used to proof the swarming behaviour, the formation of islands and the biofilm development of the two quorum-sensing knockout strains: *Δopp* and *Δhprc*. Formation of islands and biofilm images were taken as explained in previous sections.

### Phase diagram

To calculate the phase diagram of rotational Péclet number (Per) with respect to surface coverage, timelapses of swarming bacteria coming from 6 different experiments were analysed. The ‘jammed bacteria’ data plotted in the diagram (Fig. 4a) were calculated by analysing the 4 s of the time lapse prior to jam formation. Videos were acquired at 29-33 fps.

The average speed within the swarm was calculated as described earlier and used to then calculate the Pe_r_, defined as:

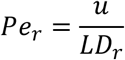

Where *u* is the average cell speed within the swarm for a given time point, *L* is the aspect ratio of the swarmers calculated to be 3.2 in average and *D_r_* the rotational diffusivity of the bacteria. The rotational diffusivity was estimated by comparing our speeds at surface coverage 0.65 with the speeds of the phase diagram in (van Damme et al., 2019) for the same surface coverage, from which D_r_ can be obtained as:

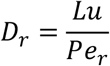

This gives *D*_r_ = 0.21 ± 0.05 s^-1^, from an average of 13 different values from 3 independent experiments. To check whether this result was sensible, we resuspended in water a sample of swarmers from a plate. Then, we recorded a video of these swarmers for 10 s and calculated D_r_ with a tracking code written in Python (Mosby et al., 2020). *D_r_* was obtained as the gradient of the angular mean-square displacement of the tracks and gave a value of D_r_ = 0.35 s^-1^. Although this is slightly higher than the previous result, it should be kept in mind that the former estimate is for cells that are moving on an agar surface, rather than swimming in a bulk fluid.

Notice that, the *Pe_r_* in Damme *et al* (van Damme et al., 2019) is obtained from instantaneous velocities of individual particles. In absence of those, we used an effective *Pe_r_* derived from the local average velocity within the swarm. This is likely to overestimate the value of *Pe*_r_ of the experimental points.

### Physical barrier

A 3% agarose solution in water was autoclaved and then poured in a Petri dish. Once it solidified, a rectangular region (6 x 1 cm) was cut out and vertically placed on a molten swarming liquid LBGM (0.5% agar). Once it solidified, swarming bacteria were inoculated in the centre of the plate. Videos of the formation of islands were recorded under 2x (Nikon Plan 2x UW, NA 0.06) in a Nikon Eclipse Ti2 microscope.

### Biofilm inhibition assay

4 *μ*l of the antibiotic kanamycin coming from four different stocks with the next concentrations: 50, 37.5, 25 and 12.5 *μ*g/*μ*l were added to four different diffusive disks. These diffusive disks were placed on a side of LBGM 0.5% agar plate for 24 h. After bacteria inoculation, the plates were incubated in a 30°C incubator for roughly 5 h until the swarming front halted. Then, additional 4 *μ*l of the next stock concentrations of kanamycin were added to the initial ones: 0, 12.5, 25 and 37.5 *μ*g/*μ*l so the total amount of kanamycin administered was kept constant and equal to 200 *μ*g.

### Wrinkle wavelength quantification

To measure the wavelength of the wrinkles we calculated the interdistance of nearest parallel wrinkles. When this was not possible, the wavelength was estimated by calculating the autocorrelation function of image intensity in space using a Fiji macro made by Michael Schmid (https://imagej.nih.gov/ij/macros/RadiallyAveragedAutocorrelation.txt) (2008) and then fitting the decay of that function to a double exponential of the form:

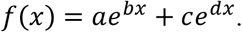

Here, the two characteristic wavelengths (1/b and 1/d) correspond to the image noise and the actual wrinkle wavelength respectively.

### Statistics

Data are reported as Mean±s.e.m. calculated from at least 3 independent experiments unless otherwise indicated.

## Acknowledgements

This research is funded by the MRC Doctoral Training Partnership (MR/N014294/1). MP and MA acknowledge support from EPSRC grant, Bridging the Gaps initiative (EP/M027503/1). MA acknowledges BBSRC/EPSRC grant to the Warwick Integrative Synthetic Biology Centre (BB/M017982/1). We thank Lewis Mosby for his help with estimates of the rotational diffusivity, and Drs. Darius Köster and Meera Unnkrishnan for their comments to the manuscript.

## References

Arlt, J., Martinez, V.A., Dawson, A., Pilizota, T., and Poon, W.C.K. (2018). Painting with light-powered bacteria. Nat. Commun. 9, 768.

Asally, M., Kittisopikul, M., Rue, P., Du, Y., Hu, Z., Cagatay, T., Robinson, A.B., Lu, H., Garcia-Ojalvo, J., and Suel, G.M. (2012). Localized cell death focuses mechanical forces during 3D patterning in a biofilm. Proc. Natl. Acad. Sci. 109, 18891–18896.

Barré, J., Chétrite, R., Muratori, M., and Peruani, F. (2015). Motility-Induced Phase Separation of Active Particles in the Presence of Velocity Alignment. J. Stat. Phys. 158, 589–600.

Bartolini, A., Tempesti, P., Ghobadi, A.F., Berti, D., Smets, J., Aouad, Y.G., and Baglioni, P. (2019). Liquid-liquid phase separation of polymeric microdomains with tunable inner morphology: Mechanistic insights and applications. J. Colloid Interface Sci. 556, 74–82.

Be’er, A., and Ariel, G. (2019). A statistical physics view of swarming bacteria. Mov. Ecol. 7.

Bechinger, C., Di Leonardo, R., Löwen, H., Reichhardt, C., Volpe, G., and Volpe, G. (2016). Active Particles in Complex and Crowded Environments. Rev. Mod. Phys. 88, 045006.

Benarroch, J.M., and Asally, M. (2020). The Microbiologist’s Guide to Membrane Potential Dynamics. Trends Microbiol. 28, 304–314.

Bhattacharyya, S., Walker, D.M., and Harshey, R.M. (2020). Necrosignaling: Cell death triggers antibiotic survival pathways in bacterial swarms. BioRxiv 2020.02.26.966986.

Butler, M.T., Wang, Q., and Harshey, R.M. (2010). Cell density and mobility protect swarming bacteria against antibiotics. Proc. Natl. Acad. Sci. 107, 3776–3781.

Cairns, L.S., Hobley, L., and Stanley-Wall, N.R. (2014). Biofilm formation by Bacillus subtilis: new insights into regulatory strategies and assembly mechanisms. Mol. Microbiol. 93, 587–598.

Cates, M.E. (2012). Diffusive transport without detailed balance in motile bacteria: does microbiology need statistical physics? Reports Prog. Phys. 75, 42601.

Cates, M.E., and Tailleur, J. (2015). Motility-Induced Phase Separation. Annu. Rev. Condens. Matter Phys. 6, 219–244.

Cates, M.E., Marenduzzo, D., Pagonabarraga, I., and Tailleur, J. (2010). Arrested phase separation in reproducing bacteria creates a generic route to pattern formation. Proc. Natl. Acad. Sci. 107, 11715–11720.

Chu, E.K., Kilic, O., Cho, H., Groisman, A., and Levchenko, A. (2018). Self-induced mechanical stress can trigger biofilm formation in uropathogenic Escherichia coli. Nat. Commun. 9.

Costerton, J.W. (1999). Bacterial Biofilms: A Common Cause of Persistent Infections. Science 284, 1318–1322.

van Damme, R., Rodenburg, J., van Roij, R., and Dijkstra, M. (2019). Interparticle torques suppress motility-induced phase separation for rodlike particles. J. Chem. Phys. 150, 164501.

Daniels, R., Vanderleyden, J., and Michiels, J. (2004). Quorum sensing and swarming migration in bacteria. FEMS Microbiol. Rev. 28, 261–289.

Digregorio, P., Levis, D., Suma, A., Cugliandolo, L.F., Gonnella, G., and Pagonabarraga, I. (2018). Full Phase Diagram of Active Brownian Disks: From Melting to Motility-Induced Phase Separation. Phys. Rev. Lett. 121, 98003.

Epstein, A.K., Pokroy, B., Seminara, A., and Aizenberg, J. (2011). Bacterial biofilm shows persistent resistance to liquid wetting and gas penetration. Proc. Natl. Acad. Sci. 108, 995–1000.

Flemming, H.-C., and Wingender, J. (2010). The biofilm matrix. Nat. Rev. Microbiol. 8, 623–633.

Frangipane, G., Dell’Arciprete, D., Petracchini, S., Maggi, C., Saglimbeni, F., Bianchi, S., Vizsnyiczai, G., Bernardini, M.L., and di Leonardo, R. (2018). Dynamic density shaping of photokinetic E. Coli. Elife 7, 1–14.

Geyer, D., Martin, D., Tailleur, J., and Bartolo, D. (2019). Freezing a Flock: Motility-Induced Phase Separation in Polar Active Liquids. Phys. Rev. X 9, 031043.

Ghosh, P., Ben-Jacob, E., and Levine, H. (2013). Modeling cell-death patterning during biofilm formation. Phys. Biol. 10, 066006.

Gonnella, G., Marenduzzo, D., Suma, A., and Tiribocchi, A. (2015). Motility-induced phase separation and coarsening in active matter. Elsevier Masson SAS 16, 316–331.

Grant, M.A.A., Waclaw, B., Allen, R.J., and Cicuta, P. (2014). The role of mechanical forces in the planar-to-bulk transition in growing Escherichia coli microcolonies. J. R. Soc. Interface 11.

Hannezo, E., and Heisenberg, C.-P. (2019). Mechanochemical Feedback Loops in Development and Disease. Cell 178, 12–25.

Hoffman, L.R., D’Argenio, D.A., MacCoss, M.J., Zhang, Z., Jones, R.A., and Miller, S.I. (2005). Aminoglycoside antibiotics induce bacterial biofilm formation. Nature 436, 1171–1175.

Jeckel, H., Jelli, E., Hartmann, R., Singh, P.K., Mok, R., Totz, J.F., Vidakovic, L., Eckhardt, B., Dunkel, J., and Drescher, K. (2019). Learning the space-time phase diagram of bacterial swarm expansion. Proc. Natl. Acad. Sci. 116, 1489–1494.

Karsenti, E. (2008). Self-organization in cell biology: A brief history. Nat. Rev. Mol. Cell Biol. 9, 255–262.

Kearns, D.B. (2010). A field guide to bacterial swarming motility. Nat. Rev. Microbiol. 8, 634–644.

Kesel, S., Grumbein, S., Tallawi, M., Marel, A., Lieleg, O., and Opitz, M. (2016). Direct Comparison of Physical Properties of Bacillus subtilis NCIB. Appl. Environ. Microbiol. 82, 2424–2432.

Krasnopeeva, E., Lo, C.J., and Pilizota, T. (2019). Single-Cell Bacterial Electrophysiology Reveals Mechanisms of Stress-Induced Damage. Biophys. J. 116, 2390–2399.

De la Fuente-Núñez, C., Reffuveille, F., Fernández, L., and Hancock, R.E.W. (2013). Bacterial biofilm development as a multicellular adaptation: Antibiotic resistance and new therapeutic strategies. Curr. Opin. Microbiol. 16, 580–589.

Lai, S., Tremblay, J., and Déziel, E. (2009). Swarming motility: A multicellular behaviour conferring antimicrobial resistance. Environ. Microbiol. 11, 126–136.

Liu, G., Patch, A., Bahar, F., Yllanes, D., Welch, R.D., Marchetti, M.C., Thutupalli, S., and Shaevitz, J.W. (2019). Self-Driven Phase Transitions Drive Myxococcus xanthus Fruiting Body Formation. Phys. Rev. Lett. 122, 248102.

Liu, J., Prindle, A., Humphries, J., Gabalda-Sagarra, M., Asally, M., Lee, D.Y.D., Ly, S., Garcia-Ojalvo, J., and Süel, G.M. (2015). Metabolic co-dependence gives rise to collective oscillations within biofilms. Nature 523, 550–554.

Lories, B., Roberfroid, S., Dieltjens, L., De Coster, D., Foster, K.R., and Steenackers, H.P. (2020). Biofilm Bacteria Use Stress Responses to Detect and Respond to Competitors. Curr. Biol. 30, 1231–1244.e4.

Lyons, N.A., and Kolter, R. (2015). On the evolution of bacterial multicellularity. Curr. Opin. Microbiol. 24, 21–28.

Mazza, M.G. (2016). The physics of biofilms - An introduction. J. Phys. D. Appl. Phys. 49.

Meredith, H.R., Srimani, J.K., Lee, A.J., Lopatkin, A.J., and You, L. (2015). Collective antibiotic tolerance: Mechanisms, dynamics and intervention. Nat. Chem. Biol. 11, 182–188.

Mosby, L.S., Polin, M., and Köster, D.V. (2020). A Python based automated tracking routine for myosin II filaments. J. Phys. D. Appl. Phys.

Nadezhdin, E., Murphy, N., Dalchau, N., Phillips, A., and Locke, J.C.W. (2020). Stochastic pulsing of gene expression enables the generation of spatial patterns in Bacillus subtilis biofilms. Nat. Commun. 11, 1–12.

Nagórska, K., Ostrowski, A., Hinc, K., Holland, I.B., and Obuchowski, M. (2010). Importance ofeps genes fromBacillus subtilis in biofilm formation and swarming. J. Appl. Genet. 51, 369–381.

Peruani, F., Klauss, T., Deutsch, A., and Voss-Boehme, A. (2011). Traffic Jams, Gliders, and Bands in the Quest for Collective Motion of Self-Propelled Particles. Phys. Rev. Lett. 106, 128101.

Preibisch, S., Saalfeld, S., and Tomancak, P. (2009). Globally optimal stitching of tiled 3D microscopic image acquisitions. Bioinformatics.

Prindle, A., Liu, J., Asally, M., Ly, S., Garcia-Ojalvo, J., and Süel, G.M. (2015). Ion channels enable electrical communication in bacterial communities. Nature 527, 59–63.

Qiu, F., Peng, G., Ginzburg, V. V, Balazs, A.C., Chen, H.-Y., and Jasnow, D. (2001). Spinodal decomposition of a binary fluid with fixed impurities. J. Chem. Phys. 115, 3779–3784.

Romero, D., Aguilar, C., Losick, R., and Kolter, R. (2010). Amyloid fibers provide structural integrity to Bacillus subtilis biofilms. Proc. Natl. Acad. Sci. 107, 2230–2234.

Schindelin, J., Arganda-Carreras, I., Frise, E., Kaynig, V., Longair, M., Pietzsch, T., Preibisch, S., Rueden, C., Saalfeld, S., Schmid, B., et al. (2012). Fiji: an open-source platform for biological-image analysis. Nat. Methods 9, 676–682.

Shemesh, M., and Chai, Y. (2013). A combination of glycerol and manganese promotes biofilm formation in Bacillus subtilis via histidine kinase KinD signaling. J. Bacteriol. 195, 2747–2754.

Su, P.T., Liao, C.T., Roan, J.R., Wang, S.H., Chiou, A., and Syu, W.J. (2012). Bacterial Colony from Two-Dimensional Division to Three-Dimensional Development. PLoS One 7, 1–10.

Sveen, J.K. (2004). An introduction to MatPIV v.1.6.1.

Vicsek, T., Czirók, A., Ben-Jacob, E., Cohen, I., and Shochet, O. (1995). Novel Type of Phase Transition in a System of Self-Driven Particles. Phys. Rev. Lett. 75, 1226–1229.

Vlamakis, H., Aguilar, C., Losick, R., and Kolter, R. (2008). Control of cell fate by the formation of an architecturally complex bacterial community. Genes Dev. 22, 945–953.

Vlamakis, H., Chai, Y., Beauregard, P., Losick, R., and Kolter, R. (2013). Sticking together: building a biofilm the Bacillus subtilis way. Nat. Rev. Microbiol. 11, 157–168.

Wang, Y., Wilks, J.C., Danhorn, T., Ramos, I., Croal, L., and Newman, D.K. (2011). Phenazine-1-carboxylic acid promotes bacterial biofilm development via ferrous iron acquisition. J. Bacteriol. 193, 3606–3617.

Wedlich-Söldner, R., and Betz, T. (2018). Self-organization: The fundament of cell biology. Philos. Trans. R. Soc. B Biol. Sci. 373.

Weitz, S., Deutsch, A., and Peruani, F. (2015). Self-propelled rods exhibit a phase-separated state characterized by the presence of active stresses and the ejection of polar clusters. Phys. Rev. E 92, 012322.

Xavier, J.B. (2011). Social interaction in synthetic and natural microbial communities. Mol. Syst. Biol. 7, 1–11.

Yan, J., Fei, C., Mao, S., Moreau, A., Wingreen, N.S., Košmrlj, A., Stone, H.A., and Bassler, B.L. (2019). Mechanical instability and interfacial energy drive biofilm morphogenesis. Elife 8, 1–28.

Yang, J., Arratia, P.E., Patteson, A.E., and Gopinath, A. (2019). Quenching active swarms: effects of light exposure on collective motility in swarming Serratia marcescens. J. R. Soc. Interface 16, 20180960.

Zhang, W., Seminara, A., Suaris, M., Brenner, M.P., Weitz, D. a, and Angelini, T.E. (2014). Nutrient depletion in Bacillus subtilis biofilms triggers matrix production. New J. Phys. 16, 015028.

